# Feedforward and feedback control share an internal model of the arm’s dynamics

**DOI:** 10.1101/362699

**Authors:** Rodrigo S. Maeda, Tyler Cluff, Paul L. Gribble, J. Andrew Pruszynski

## Abstract

Humans have a remarkable capacity to learn novel movement patterns in a wide variety of contexts. Recent work has shown that, when countering external forces, the nervous system adjusts not only voluntary (ie. feedforward) control but also reflex (ie. feedback) responses. Here we show that directly altering the physical properties of the arm (i.e. intersegmental dynamics) causes the nervous system to adjust feedforward control and that this learning also transfers to feedback responses even though they were never directly trained. In our first experiment, we altered intersegmental dynamics by asking participants to generate pure elbow movements with the shoulder joint either free to rotate or locked. Locking the shoulder joint cancels the interaction forces that arise at the shoulder during forearm rotation and thus removes the need to activate shoulder muscles to prevent shoulder joint rotation. We first asked whether the nervous system learns this altered mapping of intersegmental dynamics. In the baseline phase, we found robust activation of shoulder flexor muscles for pure elbow flexion trials prior to movement onset – as required to counter the intersegmental dynamics. After locking the shoulder joint in the adaptation phase, we found a substantial reduction in shoulder muscle activity over many trials. After unlocking the shoulder joint in the post-adaptation phase, we observed after-effects, as participants made systematic hand path errors. In our second experiment, we investigated whether such learning transfers to feedback control. Mechanical perturbations applied to the limb in the baseline phase revealed that feedback responses, like feedforward control, also appropriately countered intersegmental dynamics. In the adaptation phase, we found a substantial reduction in shoulder feedback responses – as appropriate for the altered intersegmental dynamics. We also found that this decay in shoulder feedback responses correlated across subjects with the amount of decay during feedforward control. Our work adds to the growing evidence that feedforward and feedback control share an internal model of the arm’s dynamics.

## Introduction

Humans have a remarkable capacity to modify their movements to novel situations Wolpert et al., 2011). This ability has been long studied by investigating how people change their feedforward motor commands when reaching in viscous forces fields (Shadmehr and Mussa-Ivaldi, 1994) or with altered visuomotor (Cunningham, 1989) In these cases, and in many related paradigms, the motor commands required to achieve the goal of the task are accompanied by unexpected errors and the resulting errors then induce changes in subsequent motor commands — suggesting that the nervous system updates an internal model of the environment (Wolpert et al., 1995).

Recent results suggest that such error-based learning, where participants change their feedforward motor commands to compensate for external forces that cause explicit kinematic errors, also changes how the motor system responds to sensory feedback (Wang et al., 2001; Wagner and Smith, 2008; Yousif and Diedrichsen, 2012; Cluff and Scott, 2013). For example, when participants are trained to reach in the presence of force fields and occasionally encounter experimentally-applied mechanical perturbations over the course of learning, their feedback responses to the perturbations adapt in parallel with their feedforward motor commands (Wang et al., 2001; Wagner and Smith, 2008; Yousif and Diedrichsen, 2012). Cluff and Scott (2013) and Ahmadi-Pajouh and colleagues (2012) measured muscle activity during this type of learning and showed that adapted feedback responses could be identified as early as ~50 ms following perturbation onset, suggesting that changes in feedback responses are mediated by a transcortical feedback pathway (Pruszynski and Scott, 2012; Cluff et al., 2015; Scott et al., 2015; Scott, 2016).

Here we tested whether and to what extent feedback responses adapt when people learn to compensate for novel limb dynamics that cause minimal kinematic errors. Briefly, participants moved their hand between targets placed along an arc, so that reaches could be accomplished by rotating only the elbow joint. Movements were made with the shoulder free to rotate or with the shoulder locked by the robotic manipulandum. Locking the shoulder joint physically cancels the torques that arise at the shoulder during forearm rotation (so-called interaction torques) and thus removes the need to activate shoulder muscles when making pure elbow movements. First, we tested whether people learn these altered limb dynamics during voluntary reaching movements. Since generating shoulder muscle activity does not benefit achieving the task goal and is energetically wasteful, we expected participants to reduce shoulder muscle activity following shoulder fixation. We reasoned that this learning process would be much slower than what is typically observed in paradigms where explicit movement errors occur — akin to how participants slowly optimize muscle recruitment after learning to kinematically counter a divergent force field (Franklin et al., 2004) — and perhaps explaining why previous work with similar paradigms to our own did not find learning (Koshland et al., 1991; Debicki and Gribble, 2005). Second, we examined whether learning new feedforward motor commands in this context transfers to feedback control by interspersing mechanical perturbations over the course of learning. Consistent with the hypothesis that feedforward and feedback control share an internal model of the arm’s dynamics (Wagner and Smith, 2008; Ahmadi-Pajouh et al., 2012; Cluff and Scott, 2013), we found that people learn to reduce feedforward shoulder muscle activation on a timescale of hundreds of trials and that such learning transfers to feedback control.

## Materials and Methods

### Subjects

A total of 45 healthy participants (aged 19–47, 30 females) participated in one of two experiments. All participants reported that they were right-handed and had no history of visual, neurological, or musculoskeletal impairments. Participants provided written consent, were paid for their participation, and were free to withdraw from the experiment at any time. The Office of Research Ethics at Western University approved this study.

### Apparatus

Experiments were performed using the KINARM exoskeleton robot (BKIN Technology, Kingston, ON). As previously described (Scott, 1999; Pruszynski et al., 2008, 2009), this robot permits flexion and extension movement of the shoulder and elbow joints in a horizontal plane that intersects the hand, and can independently apply torque at both joints. Visual targets and hand feedback were projected in the horizontal plane of the task via an LCD monitor and a semi-silvered mirror. Direct vision of the arm was prevented with a physical barrier. The two segments of the exoskeleton robot (upper arm and forearm) were adjusted to fit each participant’s arm and the spaces left were filled with a firm foam to ensure tight coupling with the links of the robot. The robot was then calibrated so that the projected hand cursor was aligned with each participant’s right index finger tip.

### Experiment 1: Single joint elbow reaches with shoulder fixation

Twenty participants performed twenty-degree elbow flexion and extension movements with the shoulder joint either free to move or with the shoulder fixed. At the beginning of each trial, participants moved their hand to a home target (red circle, 0.6 cm diameter). The home target position corresponded to a participant’s hand cursor position when their shoulder and elbow joints were at 10°, 60° (external angles), respectively (Figure 1, top left panel). After maintaining their hand at this location for a random period (250-500 ms, uniform distribution), a goal target (white circle: 3 cm diameter) was presented in a location that could be reached with a 20° pure elbow flexion movement. The goal target then turned red after another random period (250-500 ms, uniform distribution), cueing the participant to start their movement. At the same time, the hand feedback cursor was extinguished and remained off for the duration of the movement. Participants were instructed to move to the goal target and to do so within a specific time window. The goal target turned green when movement time (from exiting the home target to entering the goal target) was between 100 and 180 ms, orange if it was too fast (<100 ms) and red if it was too slow (>180 ms). No restrictions were placed on movement trajectories. In addition to timing constraints between targets, participants were instructed to remain at the goal target for an additional 500 ms to finish a trial. After a random period (0-1 s, uniform distribution), the goal target became a new home target (0.6 cm diameter) and the same procedure was repeated but for an extension movement.

Participants first completed 300 flexion and extension baseline trials, with the shoulder joint free to move. We then mechanically locked the shoulder joint with a physical clamp and participants repeated the same flexion and extension movements for 1100 trials (adaptation phase). Lastly, we unlocked the shoulder joint and participants again generated the same flexion and extension movements for 300 trials (post-adaptation phase) (Figure 1, top right panel). Experiment 1 lasted about 2.5 hours. Rest breaks were given throughout or when requested. Prior to data collection participants completed practice trials until they comfortably achieved ~80% success rates (approx.5 min).

**Figure 1:**
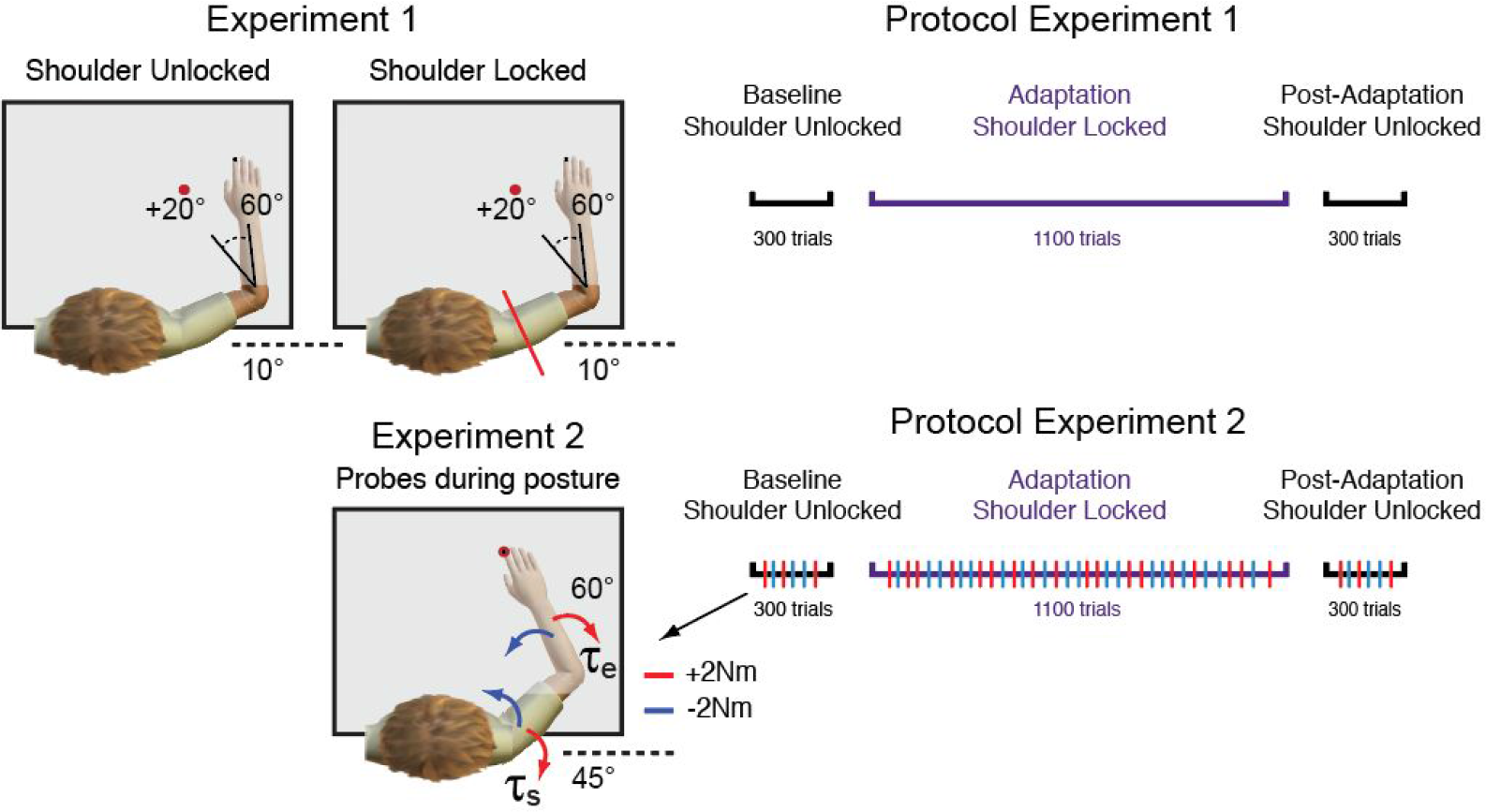
Experimental setup. In Experiments 1 and 2, participants were presented with a peripheral target that could be achieved with 20^º^ of elbow flexion rotation. Participants were instructed to perform fast and accurate reaching movements to this peripheral target and did so with their shoulder joint unlocked and locked (top left column). In Experiment 2, in addition to reaching trials, mechanical perturbations were sometimes applied (probes) to test the sensitivity of feedback responses over the course of learning (bottom left column). Red and blue arrows represent the direction of the multi-joint step-torques applied to the shoulder and elbow joints. Illustrations of the protocols for Experiments 1 and 2 are shown on the right panel. In Experiments 1 and 2, participants performed 300 baseline trials with the shoulder joint unlocked, 1100 adaptation trials with the shoulder joint locked and 300 post-adaptation trials with the shoulder joint unlocked. Multi-joint perturbations (probes, red and blue tick marks) were applied in fifteen percent of all trials in Experiments 2.

### Experiment 2: Transfer to feedback control

Twenty participants performed twenty-degree elbow flexion and extension movements with the shoulder joint free to move and with the shoulder fixed, and occasionally countered mechanical perturbations that caused pure elbow motion. As in Experiment 1, participants moved their hand to a home target (white circle, 0.6 cm diameter) to start a trial. The home target position corresponded to a participant’s fingertip when their shoulder and elbow joints were at 45° and 60° (external angles), respectively (Figure 1, bottom left panel). After a random period (250-500 s, uniform distribution), a goal target (white circle, 3cm diameter) was presented at a location that could be reached with a 20° pure elbow flexion movement. The goal target turned red after another random period (0-1 s, uniform distribution), cueing the participant to start their movement. At the same time, the hand feedback cursor was turned off and remained off for the duration of the movement. As in Experiment 1, participants were instructed to move from the home target to the goal target with a movement time ranging from 100 to 180 ms and remain in the goal target for an additional 400 ms to finish the trial. The goal target turned green when the movement was successful, orange when movement was too fast or red when it was too slow. After a random period (0-1 s, uniform distribution), the goal target became a new home target (0.6 cm diameter) and the same procedure was repeated but for an extension movement.

Participants first completed a total of 300 flexion and extension baseline trials, with the shoulder joint free to move. We then locked the shoulder joint with a servo-controller, and participants repeated the same flexion and extension movements for 1100 trials (adaptation phase). The force channel, implemented as a stiff, viscous spring and damper in the direction orthogonal to the shoulder joint (K 1000 N/m and B 250 N/(m/s)), effectively counteracted rotation of the shoulder joint (average maximum absolute deviation, ~2 deg.) and also clamped reaching trajectories. Lastly, we unlocked the shoulder joint and participants again generated the same flexion and extension movements for 300 trials (post-adaptation phase) (Figure 1, bottom right panel).

Perturbation trials occurred in fifteen percent of all trials (Figure 1, bottom left panel, probes). In these trials, when the hand cursor entered the home target, the exoskeleton gradually applied (over 2 s) a background torque of (−2 Nm/+2Nm) to the elbow to ensure baseline activation of shoulder and elbow muscles. After maintaining the cursor in the home target for a randomized duration (1.0–2.5 s, uniform distribution), a step-torque (i.e., perturbation) was applied to the shoulder and elbow joints (2 Nm at each joint over and above the background torque), which displaced the participant’s hand outside the home target. Critically, the servo-controller was turned off at perturbation onset and we chose this combination of shoulder and elbow loads to minimize shoulder motion (see Kurtzer et al., 2008; Maeda et al., 2017). Participants were instructed to quickly counter the load and bring their hand back to the goal target (centered on the home target). If the participant returned to the goal target within 385 ms of perturbation onset, the target circle changed from white to green, otherwise the target circle changed from white to red. In five percent of all trials, the background torques turned on, remained on for the same time period (1.0–2.5 s, uniform distribution), but then slowly turned off, after which participants were still required to perform the reaching movements. These trials ensured that background loads were not always predictive of perturbation trials.

The order of all perturbation, control and reaching trials was randomized in the baseline and post adaptation phases and pseudo randomized in blocks of every 22 trials in the adaptation phase. Experiment 2 lasted about 2.5h. Rest breaks were given throughout or when requested. Prior to data collection participants completed practice trials until they comfortably achieved ~80% success rates (approx. 5 min).

Five additional participants performed the same version of this Experiment 2 without locking the shoulder joint. This served as a control for both Experiment 1 and 2 to rule out changes in feedforward or feedback changes caused by extensive practice rather than the shoulder locking manipulation.

### Kinematic recordings and analysis

Movement kinematics (i.e. hand position, and joint angles) were sampled at 1000Hz and then low-pass filtered (12 Hz, 2-pass, 4th-order Butterworth). In Experiment 1, all data were aligned on movement onset in Experiment 1. In Experiment 2, data from reaching trials was aligned on movement onset and data from perturbation trials was aligned on perturbation onset. Movement onset was defined as 5% of peak angular velocity of the elbow joint (see Gribble and Ostry, 1999; Maeda et al., 2017). We quantified the adaptation and aftereffects of reaching movements following shoulder fixation using hand path errors relative to the center of the target at 80% of the movement between movement onset and offset (also defined at 5% of peak angular velocity of the elbow joint). This corresponds to 170 ms (SD 15 ms) duration. This window was chosen to select the kinematic traces before any feedback corrections.

### EMG recordings and analysis

We measured electromyographic signals from upper limb muscles using surface electrodes (Delsys Bagnoli-8 system with DE-2.1 sensors, Boston, MA). Electrodes were placed on the skin surface overlying the belly of five muscles (pectoralis major clavicular head, PEC, shoulder flexor; posterior deltoid, PD, shoulder extensor; biceps brachii long head, BB, shoulder and elbow flexor, Brachioradialis, BR, elbow flexor; triceps brachii lateral head, TR, elbow extensor). Before electrode placement, the participants’ skin was abraded with rubbing alcohol, and the electrodes were coated with conductive gel. Electrodes were placed along the orientation of muscle fibers. A reference electrode was placed on the participant’s left clavicle. EMG signals were amplified (gain = 10^3^), and then digitally sampled at 1,000 Hz. EMG data were then band-pass filtered (20–500 Hz, 2-pass, 2nd-order Butterworth) and full-wave rectified.

In Experiment 1, we investigated whether shoulder muscles adapt to novel intersegmental dynamics following shoulder fixation. To compare the changes in amplitude of muscle activity over time and across different phases of the protocol, we calculated the mean amplitude of phasic muscle activity across a fixed time-window, −100 ms to +100 ms relative to movement onset, as has been done previously (see Debicki and Gribble, 2005; Maeda et al., 2017). These windows were chosen to capture the agonist burst of EMG activity in each of the experiments, but our results did not qualitatively change with small changes in this averaging window.

In Experiment 2, we investigated whether feedback responses in shoulder muscles also adapt to the novel intersegmental dynamics following shoulder fixation. To test whether the short and long latency stretch response of shoulder flexors account for and adapt to novel intersegmental dynamics, we binned the PEC EMG into previously defined epochs(see Pruszynski et al., 2008). This included a pre-perturbation epoch (PRE, −50-0 ms relative to perturbation onset), the short-latency stretch response (R1, 25-50 ms), the long-latency stretch response (R2/3, 50-100 ms), and the voluntary response (VOL, 100-150ms).

Normalization trials prior to each experiment were used to normalize muscle activity such that a value of 1 represents a given muscle sample’s mean activity when countering a constant 1 Nm torque (see Pruszynski et al., 2008; Maeda et al., 2017). Data processing was performed using MATLAB (r2016b, Mathworks, Natick, MA). For simplicity, here we only report the results of flexion movements and for feedback responses in shoulder flexors, however the results are similar for the extension.

### Statistical analysis

All statistical analyses were performed using R (RStudio, Boston, MA). We performed different statistical tests (e.g., repeated measures ANOVA with Tukey tests for multiple comparisons, t-test, and regression analysis), when appropriate in each of the two experiments. Details of these procedures are provided in the Results. Experimental results were considered statistically significant if the corrected p-value was less than < 0.05.

## Results

### Experiment 1: Single-joint elbow reaching with shoulder fixation

Participants (N = 20) had no difficulty learning the imposed speed and accuracy constraints and achieved >90% success within 5 minutes of practice. Although never instructed to do so, even with the shoulder free to move participants moved from the start target to the goal target by almost exclusively rotating their elbow joint (Figure 2A). Despite minimal shoulder rotation, we found substantial shoulder flexor muscle activity prior to movement onset, as required to compensate for the torques that arise at the shoulder when the forearm rotates (Gribble and Ostry, 1999; Maeda et al., 2017) (Figure 2B).

After the baseline trials, we physically locked the shoulder joint of the KINARM, a manipulation which eliminated all the torques that needed to be countered by muscles spanning the shoulder joint when the forearm rotated. Locking the shoulder joint did not alter task performance, with participants continuing to demonstrate >90% success rates.

**Figure 2:**
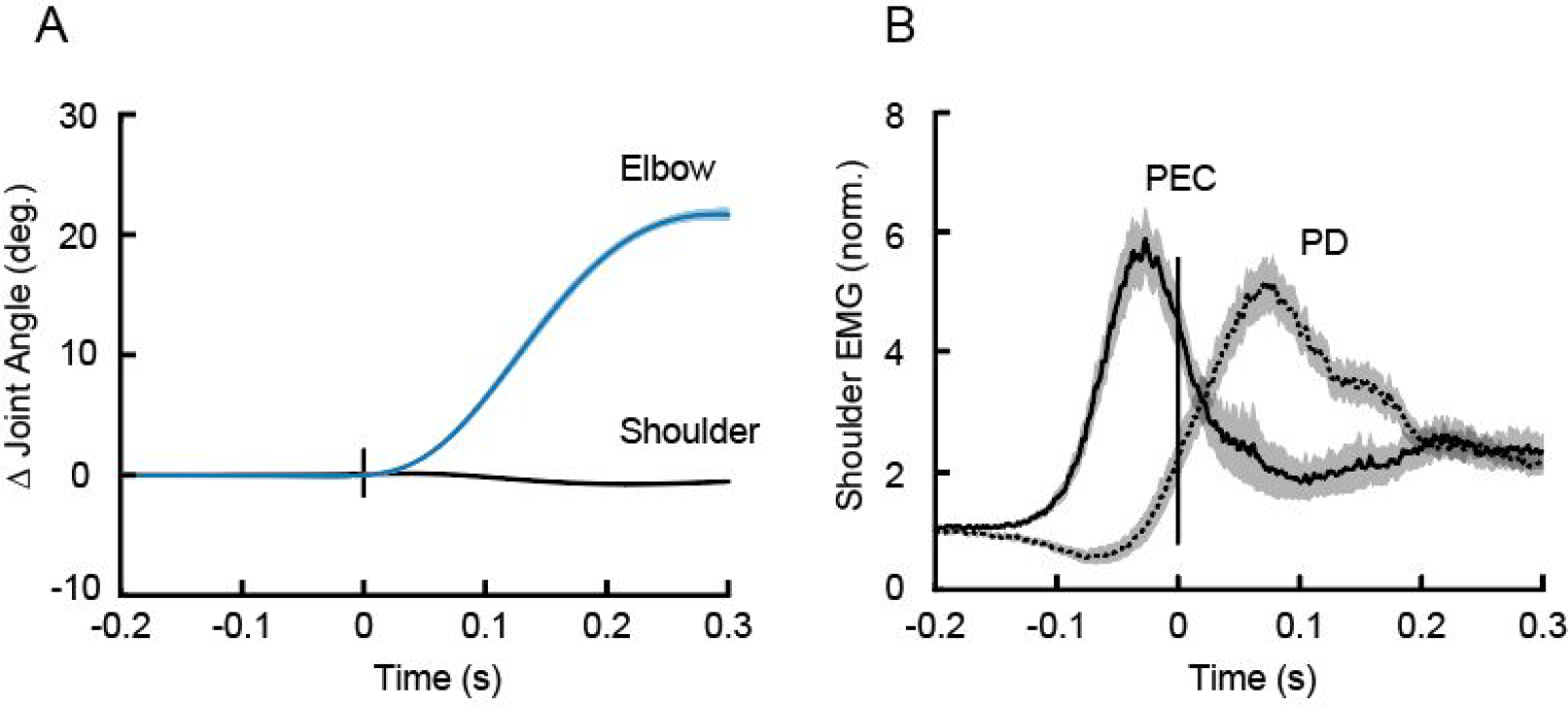
Compensating for intersegmental dynamics during self-initiated elbow reaches. (A) Average kinematics of the shoulder (black) and elbow (blue) joints for elbow flexion movements with the shoulder joint free to move (i.e. baseline trials). Shaded areas represent the standard error of the mean. Data are aligned on movement onset. (B) Solid and dashed lines represent average agonist (PEC) and antagonist (PD) muscle activity during flexion movements, respectively. EMG data are normalized as described in the Methods. Data are aligned on movement onset.

We first tested whether the nervous system adapts to the experimentally imposed intersegmental dynamics by reducing shoulder muscle activity. Figure 3A illustrates mean shoulder flexor muscle activity, in a fixed time window (−100 to 100 ms, see Methods) relative to movement onset, across trials both before (i.e. baseline epoch) and after (i.e. adaptation epoch) the shoulder was physically locked. The magnitude of shoulder muscle activity appeared to slowly decrease over the course of the adaptation trials. We then removed the shoulder lock and found that shoulder muscle activity quickly returned to baseline levels (post-adaptation trials in Fig. 3A). Indeed, a one-way ANOVA comparing shoulder flexor muscle activity late (last 25 trials) in the baseline, adaptation and post-adaptation phases revealed a reliable effect of phase on shoulder muscle activity (F2_2,38_ = 17.103, P < 0.0001). Tukey post-hoc tests showed that shoulder flexor muscle activity reliably decreased by 36% relative to baseline (P = 0.0001) when the shoulder was locked and then reliably increased again after the shoulder was unlocked, returning to that seen in baseline trials (P = 0.5; Figure 3B,C). Note that we found no corresponding changes in elbow muscle activity as a function of epoch (one-way-ANOVA, F_2,36_ = 1.505, P = 0.236; Figure 3D-F). Moreover, in a control experiment where participants (N = 5) performed the same number of total trials but never with the shoulder locked, we found no reliable decrease in shoulder muscle activity over trials corresponding to the phases of the main experiment (F_2,8_ = 1.307 P = 0.323; see gray error bars in Fig. 3A). Indeed, shoulder muscle activity at the end of the adaptation phase in the main experiment was reliably smaller than at the equivalent point in the control experiment (t_19_ −3.7, P = 0.001), indicating that shoulder muscle activity decay is related to shoulder fixation.

**Figure 3:**
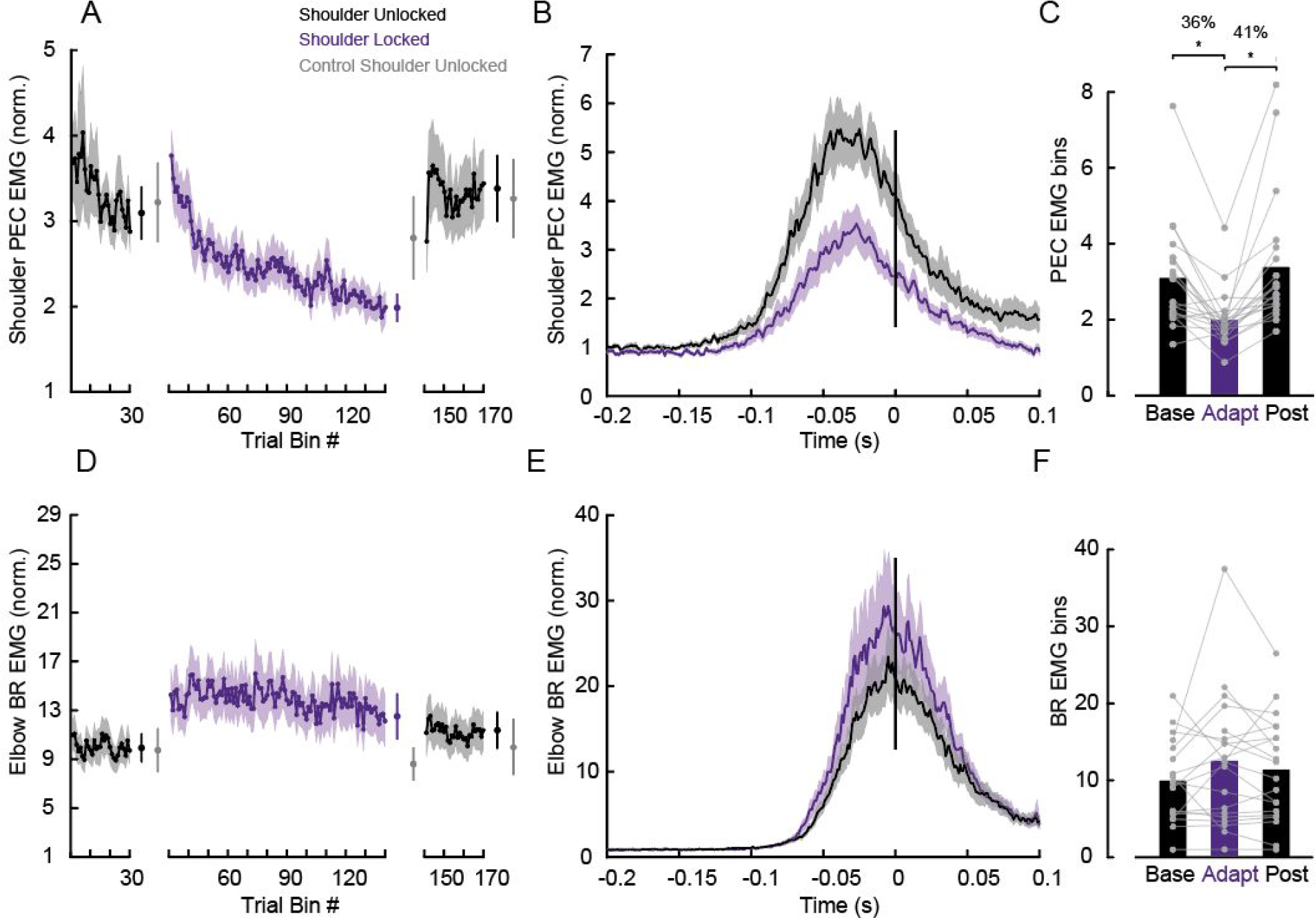
Learning novel intersegmental dynamics following shoulder fixation. (A) Average PEC muscle activity in a fixed time window (−100 to 100 ms relative to movement onset). Each data bin is 5 trials. Shaded areas represent the standard error of the mean. EMG data are normalized as described in the Methods. Error bars plotted between the epochs represent the mean and standard error of the last 5 bins of trials in each phase contrasted with the respective bins in the control experiment in gray. (B) Time series of PEC normalized muscle activity averaged over the last 25 baseline and adaptation trials. Data is aligned on movement onset. (C) Average PEC muscle activity in a fixed time window (−100 to 100 ms relative to movement onset) associated with these trials. Each dot represents data from a single participant. Asterisks indicate reliable effects (p < 0.05, see main text). (D-F) data for elbow BR muscle are shown using the same format as A-C.

In addition to reduced shoulder muscle activity, another indication that participants learned the novel limb dynamics we imposed was the presence of robust after-effects. That is, in the early post-adaptation phase, participants produced substantial reaching errors in the direction that one would predict if they failed to compensate for the unlocked intersegmental dynamics (Figure 4A). Unlike the introduction of the shoulder lock, the kinematic after-effects present when the shoulder was unlocked resulted in participants not achieving the speed and accuracy constraints imposed in our task – that is, participants made errors when the shoulder was unlocked. We performed a one-way ANOVA to compare reach accuracy (measured as distance from the center of the goal target) of trials late in the baseline phase (last 25 trials), trials early in the post-adaptation phase (first 3 trials) and trials late (last 25 trials) in the post-adaptation phase (Figure 4B). Note that we chose a smaller bin size early in the post adaptation because the return to baseline after unlocking the shoulder joint happens quickly (Figure 3A). We found a significant effect of phase (F_2,38_ = 4.65, P = 0.01). Tukey post-hoc tests showed that movement errors increased by 42% (p=0.03) from baseline to early post-adaptation and returned to baseline levels (p=0.97) in late post-adaptation trials.

**Figure 4:**
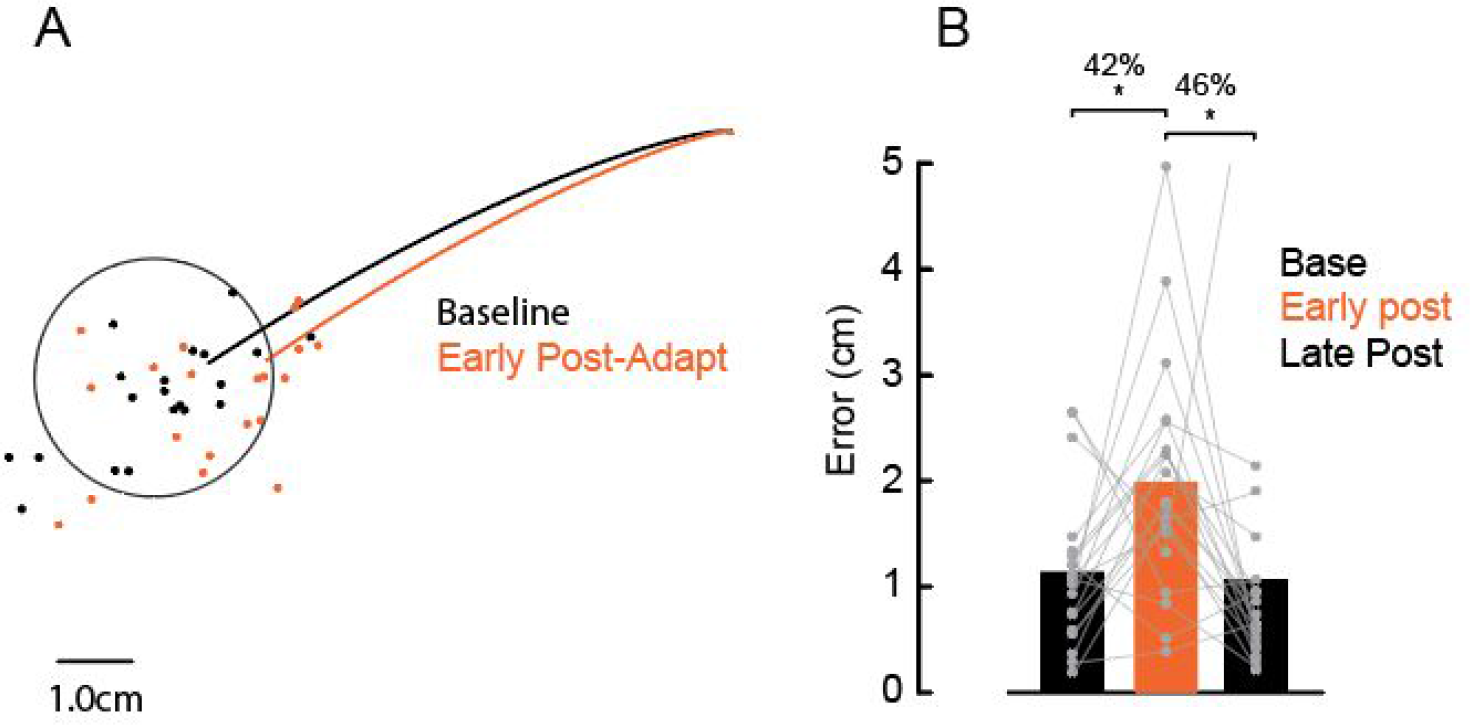
Movement trajectories after adapting elbow reaches. (A) Average hand trajectories late in the baseline (25 trials) and early in the adaptation trials (first 3 trials). Each dot represents data from a single participant. (B) Average error between hand position at movement offset to the center of the target in the last 25 trials in the baseline, first 3 trials early in the post-adaptation and last 25 trials late in post-adaptation phases (p < 0.05, see main text).

### Experiment 2: Transfer to feedback control

Participants (N = 20) again moved their hand between two targets that required 20 degrees of elbow flexion (Figure 5A). Participants had no difficulty with the imposed speed and accuracy constraints of the task and achieved >90% of success within 5 min of practice. In addition to these reaching trials, we occasionally applied mechanical perturbations to the arm while participants maintained their hand in the home target. The mechanical perturbations consisted of step torques applied simultaneously to the shoulder and elbow chosen so they caused minimal shoulder motion but different amounts of elbow motion (Figure 5B).

As shown in Experiment 1 (Figure 2) and as previously demonstrated, participants completed the reaching trials almost exclusively by rotating their elbow joint and, in so doing, generated a substantial amount of shoulder muscle activity (Figure 5C) (Gribble and Ostry, 1999; Maeda et al., 2017). Also, as previously demonstrated, we found that mechanical perturbations that created pure elbow motion elicited substantial shoulder muscle activity in the long-latency epoch (Kurtzer et al., 2008; Maeda et al., 2017) (Figure 5D), as appropriate for countering the imposed joint torques.

**Figure 5:**
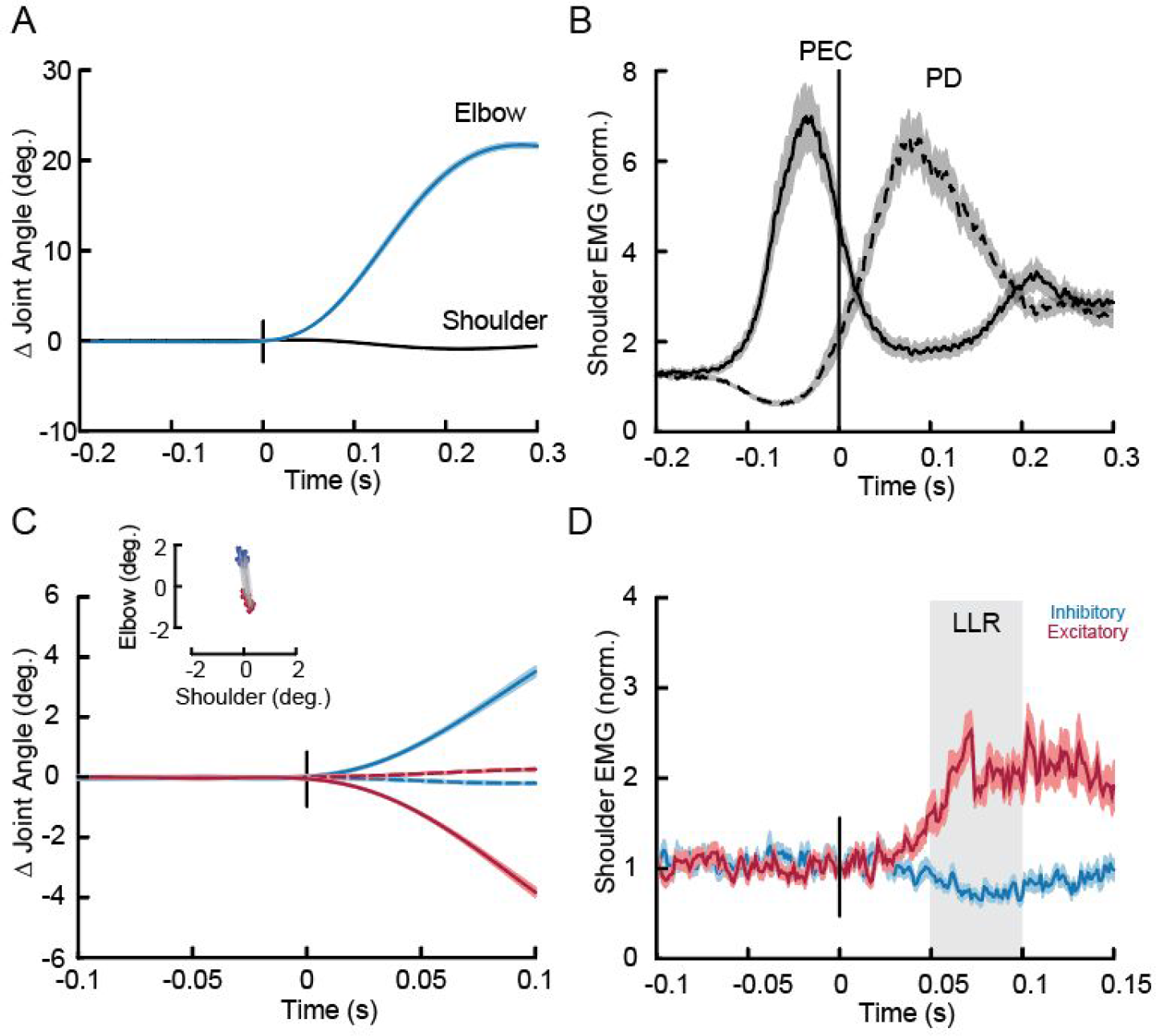
Compensating for intersegmental dynamics during self-initiated reaches and when responding to perturbations. (A) Average kinematics of the shoulder (dashed) and elbow (solid) joints for elbow flexion trials. Shaded areas represent the standard error of the mean. Data are aligned on movement onset. (B) Dashed and solid lines represent average agonist (PEC) and antagonist (PD) muscle activity associated with the movement in A. Data are aligned on movement onset. (C) Average kinematics of the shoulder (dashed) and elbow (solid) joints following mechanical perturbations. Red and blue traces are from the shoulder/elbow extensor torque and shoulder/elbow flexor torque conditions, respectively. Inset shows the amount of shoulder and elbow displacement 50 ms post-perturbation (data are shown for all subjects). (D) Normalized shoulder muscle activity associated with panel C.

We then tested whether, as in Experiment 1, feedforward commands to shoulder muscles adapt when participants produce the same elbow rotations following shoulder fixation. Consistent with Experiment 1, we found a reliable effect of phase on shoulder muscle activity (F_2,38_ = 19.7 P < 0.0001) with Tukey post-hoc tests showing that PEC muscle activity decreased by 41% (P=0.0001) following shoulder fixation and increased when removing the shoulder clamp by 31% (P=0.001). We again found no corresponding changes in elbow muscle activity as a function of epoch (one-way-ANOVA, F_2,38_ = 0.075, P = 2.761; supplemental Figure S1 C-D). Participants again showed evidence of after-effects in the early post-adaptation phase, producing substantial reaching errors in the direction required to compensate for intersegmental dynamics (F2,38 = 14.4, P < 0.001) with Tukey post-hoc tests revealing that movement errors increased by 50% (P<0.001) from baseline to early post-adaptation and returned to baseline levels (P<0.61) in late post-adaptation trials (see supplemental Figure S1E-F).

The main goal of Experiment 2 was to examine whether learning novel intersegmental dynamics following shoulder fixation during voluntary control also modifies the sensitivity of sensory feedback responses to mechanical perturbations. We tested this idea by occasionally applying mechanical perturbations across all phases of the protocol. If the shoulder joint was locked, we unlocked the shoulder joint and then applied mechanical perturbations at the shoulder and elbow joints that created pure elbow motion. Thus, all joints were free to rotate in perturbation trials across all phases of the protocol such that participants only experienced altered intersegmental dynamics in voluntary reaching trials.

**Figure 6:**
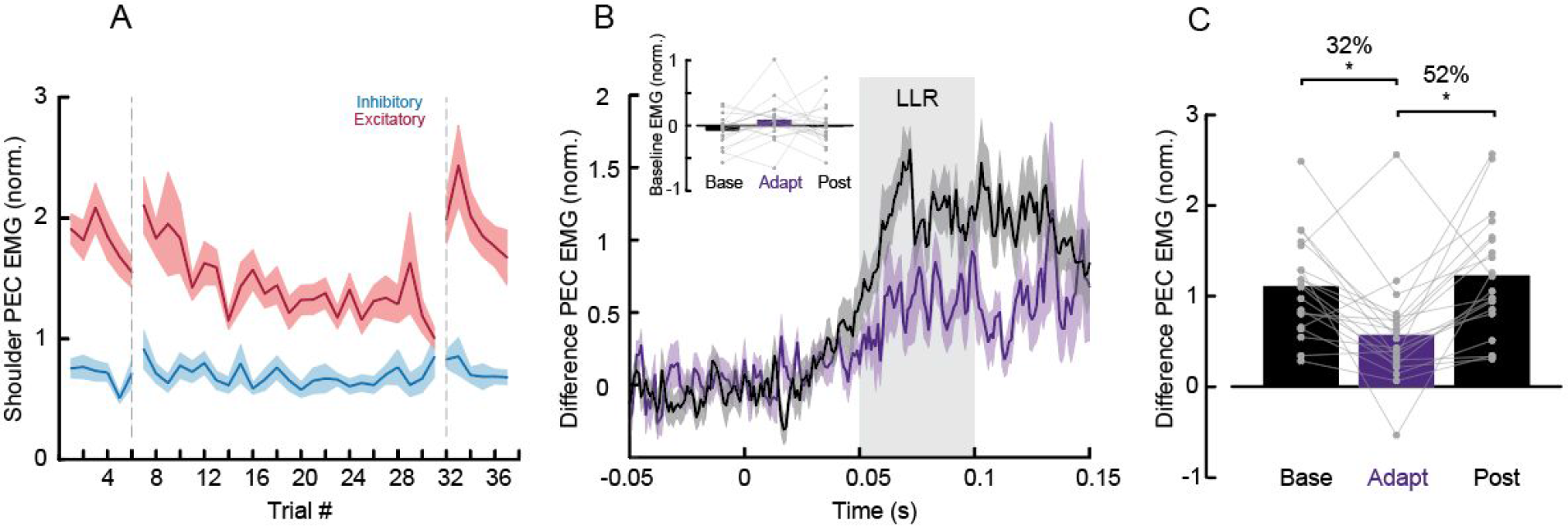
Rapid feedback responses following learning during self-initiated reaching with shoulder fixation. (A) Average shoulder EMG (filtered and rectified) in the long latency epoch (50-100ms) across trials. Red and blue traces indicate the shoulder/elbow extensor torque (excitatory), and shoulder/elbow flexor torque conditions (inhibitory), respectively. Vertical dashed line separates perturbation trials that happened in the baseline, adaptation and post-adaptation phases. (B) 5 perturbation trials of the difference of PEC muscle activity (excitatory-inhibitory) averaged in late baseline and late adaptation trials. Baseline pre-activity in these trials is shown in the inset. (C) Long latency epoch of the difference of PEC muscle activity (excitatory-inhibitory) in the baseline, adaptation and post-adaptation phases. (p < 0.05, see main text).

Figure 6A illustrates group mean shoulder (PEC) muscle activity in the long-latency epoch over trials in the baseline, adaptation and post-adaptation phases of the protocol. Red and blue traces indicate the shoulder/elbow extensor torque, and shoulder/elbow flexor torque conditions (PEC EMG excitatory and inhibitory loads). We first took the difference between excitatory and inhibitory traces as a metric of sensitivity to intersegmental dynamics. We then used a one-way ANOVA to compare this difference of PEC muscle activity in the long-latency epochs, averaged in last 5 trials of the baseline, adaptation and post-adaptation phases. We found a reliable effect of phase (F_2,38_ = 9.851, P< 0.0001). Consistent with our prediction, Tukey post-hoc tests showed that the difference in PEC muscle activity in the long-latency epoch decreased by 48% (p=0.001) following shoulder fixation and returned to baseline levels in the post-adaptation phase (p=0.73) (Figure 6B-C). We performed the same analysis to assess for changes also in short-latency epoch, but we found no reliable differences (F_2,38_ = 0.236, P<0.791). Importantly, we tested whether there was a change in baseline EMG activity pre-perturbation across phases, which could potentially explain these changes in EMG in the long-latency epoch (gain scaling, Pruszynski et al., 2009). We used a one-way ANOVA to compare the baseline activity of PEC muscle activity pre-perturbation as a function of experimental phase and found no reliable effect (F_2,38_ = 1.649 P = 0.206; Figure 6B, inset). Moreover, in a control experiment where participants (N = 5) performed the same number of total trials but never with the shoulder locked, we found no decrease in the long-latency epoch over trials corresponding to the phases of the main experiment (F_2,8_ = 0.449 P = 0.653).

Finally, we tested whether there was a correlation between learning effects associated with feedforward motor commands and feedback responses. We found a reliable linear relationship between the decrease in shoulder muscle activity during voluntary elbow rotations while the KINARM shoulder joint is fixed, and the decrease in shoulder muscle activity measured in the long-latency epoch of perturbation trials (Slope= 0.45, R=0.47, p=0.03) (Figure 7).

**Figure 7:**
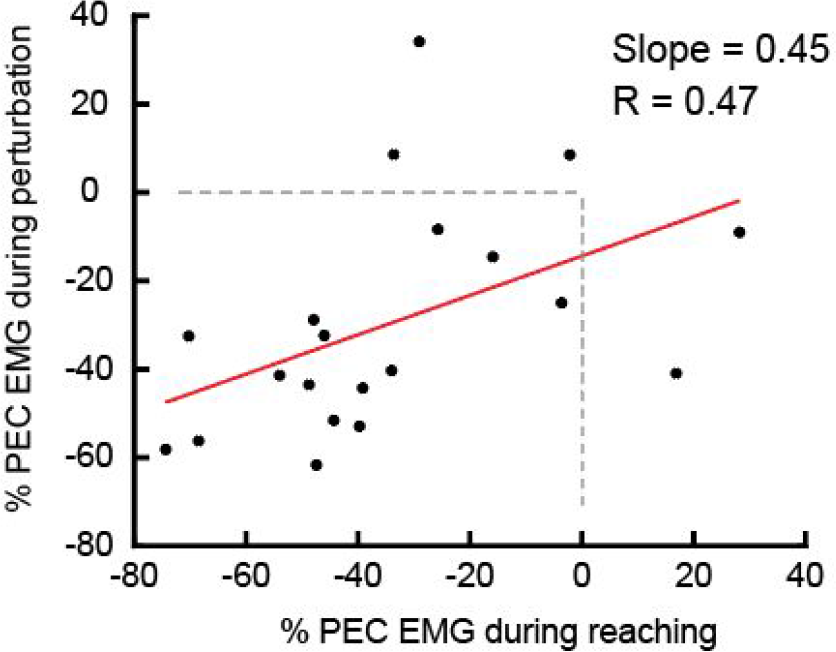
Correlates of reach adaptation and feedback responses. (A) Vertical axis is percent change in the long-latency epoch responses (50-100 ms) between baseline and late adaptation phase. Horizontal axis is percent change in muscle activity during elbow reaching trials (−100 to 100 ms relative to movement onset).

## Discussion

In Experiment 1, we tested whether the nervous system learns novel intersegmental dynamics during feedforward control. Consistent with this idea, we found that agonist shoulder muscle activity during voluntary elbow rotations slowly decreased after the shoulder was mechanically fixed. We also found systematic trajectory errors (i.e. after-effects) after removing shoulder fixation. In Experiment 2, we tested whether learning novel intersegmental dynamics during feedforward control transfers to feedback control. Consistent with this idea, we found that shoulder muscle activity in the long-latency stretch epoch slowly decreased after the shoulder was mechanically fixed, even though the long-latency stretch response was never directly exposed to the novel intersegmental dynamics. Taken together, our results demonstrate that the nervous system learns novel intersegmental dynamics following shoulder fixation, and that this learning transfers from feedforward to feedback control.

### Learning novel intersegmental limb dynamics during feedforward control

Intersegmental dynamics complicates the mapping between joint torques and joint motions. One way to study how the nervous system learns and compensates for intersegmental dynamics is by experimentally manipulating these rotational forces during movement. For example, Sainburg and colleagues (1999) investigated whether the nervous system learns novel intersegmental dynamics in an error-based paradigm by changing the center of mass of the forearm and thus altering intersegmental dynamics during reaching. Consistent with many other error-based learning paradigms (Wolpert et al., 2011), reaching was initially altered, but participants were able to correct their movement trajectories over a relatively small number of trials, and they showed large trajectory errors after removing the added mass (ie. after-effects). Debicki and Gribble (2005) took another approach. They explored how the nervous system learns novel intersegmental dynamics by asking participants to generate pure elbow flexion movements with the shoulder joint free to move or mechanically fixed. Fixing the shoulder joint removes the interaction torques that arise at the shoulder due to forearm rotation and thus removes the need to activate shoulder muscles. In this paradigm, participants don’t make substantial kinematic errors when the shoulder is fixed so reducing shoulder muscle activity is not strictly necessary to achieve the task. Indeed, they report that shoulder muscle activity remains unchanged after it is fixed. Wrist muscle activity also appears to remain unchanged when the wrist joint is fixed during elbow rotation (Koshland et al., 1991). Using the same paradigm, we show that such learning does happen: shoulder muscle activity decreases and people show kinematic after–effects. However, learning unfolds over a timescale much longer than previously examined and what is typical for error-based learning experiments. Even over 550 trials learning was incomplete and shoulder muscle activity did not reach steady-state.

Why was learning incomplete? The most likely possibility is that learning new intersegmental dynamics simply takes place over much longer time scale (e.g. over development) so we did not observe complete learning even with our extended protocol. Indeed, in our results, learning did not reach a clear steady state suggesting learning was still ongoing (see Figure 3A). Alternatively, the burst of shoulder muscle activity that is observed just prior to elbow rotation may partially reflect a hard-wired synergy between shoulder and elbow muscles that makes a complete dissociation impossible (Koshland et al., 1991; Debicki and Gribble, 2005; de Rugy et al., 2012). Insight into these possibilities will ultimately come from long-term learning studies, either by having participants come back over many sessions and exploiting extended periods of practice or by looking at muscle coordination in people who have their joints immobilized for long periods of time (e.g. after bone fracture).

Our finding that learning without kinematic errors is slow is consistent with other studies examining how motor learning, in the context of reaching, evolves in the absence of kinematic errors (Diedrichsen et al., 2010; Vaswani and Shadmehr, 2013). For example, Vaswani and Shadmehr (2013) used a force channel to clamp trajectory errors to zero after participants had learned a force field and showed that memories of the force field fade on a time scale much longer than the original learning. Others have emphasized that motor learning continues to optimize motor programs after kinematic errors have been eliminated, presumably to make the movement more efficient and reduce metabolic cost (Takahashi et al., 2006; Emken et al., 2007; Huang and Ahmed, 2014). Although learning without kinematic errors usually happens slowly, this is not always the case. For example, Cordo and Nashner (1982) asked participants to push and pull a handle while standing freely and with their trunk stabilized, quite analogous to our own study. They found a reduction in automatic postural responses after only a few trials of trunk stabilization. This may be a special case where the context is very explicit and the switch well practiced (as people often have their trunk stabilized when sitting), and where the potential energy savings associated with not activating core muscles is relatively high.

### Learning novel intersegmental limb dynamics during feedforward control transfers to feedback control

There is growing evidence that motor learning can modify how the motor system responds to sensory feedback (Wang et al., 2001; Kimura et al., 2006; Wagner and Smith, 2008; Ahmadi-Pajouh et al., 2012; Yousif and Diedrichsen, 2012; Cluff and Scott, 2013). In previous studies, participants made reaching actions in the presence of force fields that caused kinematic errors and experienced occasional mechanical perturbations to probe feedback responses over the course of learning. These studies convincingly show that, when feedforward motor commands adapt, feedback responses to mechanical perturbations also adapt. Our results reveal that the same is true when people learn new intersegmental dynamics in the absence of systematic kinematic errors. Since we ensured that fast feedback responses were never exposed to the learning context, the transfer we report shows that feedforward and feedback responses have access to a shared internal model of the arm’s dynamics.

Internal models are a central concept in motor control, including actions like reaching, grasping and object manipulation (Wolpert et al., 2011). They enable the nervous system to predict the consequences of the motor commands it generates and to determine which motor commands are required to execute a particular action, critical computations for stable and accurate control given various sources of noise and delays in the sensorimotor system (Wolpert et al., 1995; Harris and Wolpert, 1998). Although internal models have been extensively studied in the context of feedforward motor commands, feedback responses also rely on an internal model that has many of the key features of feedforward motor commands (Lacquaniti and Soechting, 1986; Kurtzer et al., 2008, 2009, 2014; Pruszynski et al., 2011; Crevecoeur and Scott, 2013, Soechting, 1986; Kurtzer et al., 2008, 2009, 2014; Pruszynski et al., 2011; Crevecoeur and Scott, 2013, 2014). For example, long-latency stretch responses respond to the expected future kinematic state of the arm by integrating incoming sensory information with prior knowledge about the mechanical perturbations encountered in the environment (Crevecoeur and Scott, 2013).

To our knowledge, no studies have directly addressed the neural mechanisms that underlie shared internal models for feedforward and feedback control and this is an important gap in the literature. However, some of the same neural structures have been implicated in housing internal models for feedforward or feedback control in isolation (For review, see Kurtzer, 2014). One likely locus is the primary motor cortex (M1). In terms of feedforward control, Gritsenko et al. (2011) used transcranial magnetic stimulation applied to human primary motor cortex while participants reached to targets placed at locations that yielded assistive or resistive interaction torques between the arm and forearm. Their results showed that motor evoked potentials were greater for movement directions that included resistive interaction torques compared to assistive movements, indicating that M1 mediates feedforward compensation for the arm’s intersegmental dynamics. In terms of feedback control, Pruszynski et al. (2011) showed that transcranial magnetic stimulation applied to human M1 potentiates shoulder muscle responses following mechanical perturbations that cause pure elbow displacement, indicating that M1 mediates feedback compensation for the arm’s intersegmental dynamics. In fact, many of the single neurons in monkey M1 that are activated during feedforward generation of motor commands also respond to mechanical perturbations (Evarts, 1973; Evarts and Tanji, 1976; Wolpaw, 1980; Evarts and Fromm, 1981; Picard and Smith, 1992; Herter et al., 2009 Pruszynski et al., 2011; Omrani et al., 2014; Pruszynski, 2014). Although it is unknown whether and how these specific neurons modify their responses in the context of motor learning, primary motor cortex is intimately involved in motor learning so such overlap is plausible if not likely (for review, see Sanes and Donoghue, 2000; see also Kawai et al., 2015). Another likely locus is the cerebellum, which, at the highest level, is thought to contain the internal models that underlie voluntary motor control (Wolpert et al., 1998) Consistent with this role, damage to the cerebellum yields profound deficits coordinating the joints without affecting the ability to generate the required forces (Holmes, 1939; Goodkin et al., 1993; Bastian et al., 1996, 2000). Cerebellum also contributes to feedback control. Neurons in the dentate and interpositus nuclei of the cerebellum rapidly respond to mechanical perturbations in a goal–dependent manner (Strick, 1983) and long-latency stretch responses are reduced in patients with cerebellar dysfunction (Hore and Vilis, 1984; Kurtzer et al., 2013). Of course, these cerebellum and primary motor cortex do not act in isolation and an important line of future research is precisely delineating their interactions along with other cortical and brainstem contributors.

---

**Figure S1:**
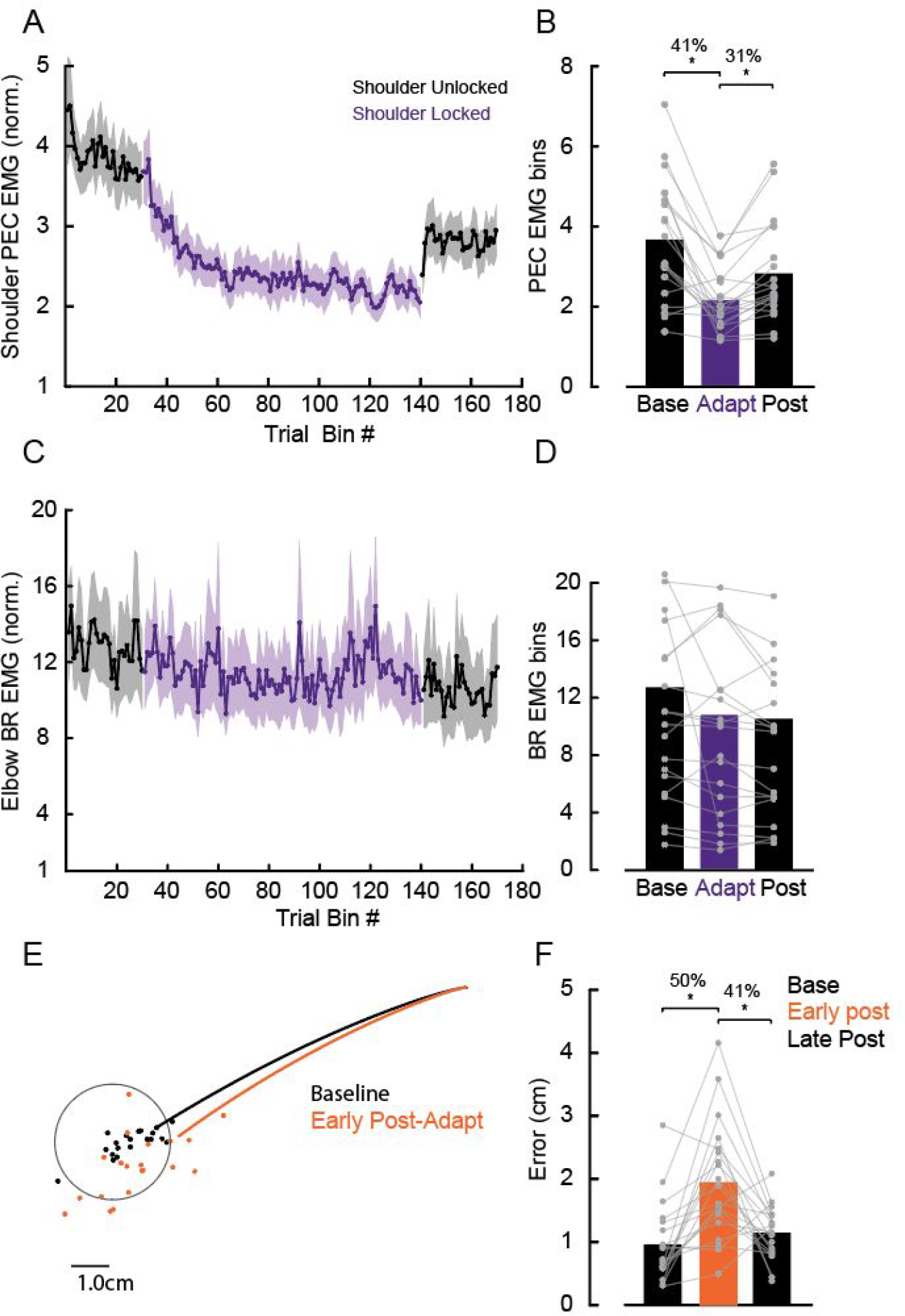
Learning novel intersegmental dynamics following shoulder fixation (Experiment 2). (A) Mean PEC muscle activity in a fixed time window over trials (−100 to 100 ms relative to movement onset). Shaded areas represent the standard error of the mean. (B) Mean PEC muscle activity in a fixed time window (−100 to 100 ms relative to movement onset) of 25 trials late in the baseline, adaptation and post-adaptation phases. (C-D) data for elbow BR muscle activity in the same format as shown in (A-B). (E) Mean hand trajectories in the last 25 trials in the baseline and first 3 post-adaptation trials. Each dot represents data from a single participant at 80% of reach. (F) Mean error between hand position to the center of the target associated with these trials in the baseline, early and late in the late post-adaptation phases. Each dot represents data from a single participant. Asterisks indicate reliable effects (p < 0.05, see main text).

